# COBRA: Cell-type-specific Orthogonal Batch effect Removal Algorithm in single cell RNA-sequencing data

**DOI:** 10.64898/2025.12.04.692478

**Authors:** Sujin Seo, Sungho Won, Kyungtaek Park

**Author notes:** **Corresponding author:** Kyungtaek Park, Ph.D., Department of Statistics, Jeonbuk National University, Jeonju, Republic of Korea, Sungho Won, Ph.D., Department of Public Health Sciences, Seoul National University, Seoul, Republic of Korea. These authors contributed equally to this work (Sungho Won, Kyungtaek Park).

## Abstract

Single-cell RNA sequencing (scRNA-seq) enables high-resolution profiling of cellular heterogeneity, yet batch effects remain a critical challenge in data integration. Existing batch correction methods often assume homogeneous batch effect across cell types, operate in reduced-dimensional space leading to potential loss of biological information, and require extensive computational resources. Here, we introduce COBRA (https://github.com/wonlab-healthstat/COBRA), a linear model-based batch correction method that explicitly adjusts cell-type-specific batch effect. By orthogonalizing batch-associated parameters with respect to biological variables, COBRA removes technical artifacts while preserving biologically meaningful transcriptional differences. When cell type annotations are unavailable, COBRA implements an iterative clustering algorithm to estimate pseudo-cell types while accounting for batch effects. COBRA retains the full gene expression matrix, ensuring seamless integration for downstream analyses. We evaluated COBRA across simulated and real-world datasets, including type 2 diabetes, and COVID-19 datasets. COBRA outperformed in terms of batch mixing efficiency, preservation of biological group structure, and accuracy of differentially expressed gene detection.

## Introduction

Single-cell RNA sequencing (scRNA-seq) has revolutionized our ability to explore cellular heterogeneity and gene expression profiles at the single cell level. As the number of publicly available scRNA-seq datasets continues to grow, integration of multiple datasets has become essential for comprehensive analysis. However, technical variations often arise due to differences in sample preparation, sequencing platforms, and data processing steps across batches (1). Without proper correction of these nuisance variations, misleading conclusions may be drawn (2). Moreover, inherent biological heterogeneity contributes to the complexity of scRNA-seq data, requiring sophisticated methods to accurately remove the non-biological effects while preserving true biological signals (3). The high dimensionality and the diverse biological responses across different conditions necessitate computationally efficient and robust correction methods.

To address these challenges, a variety of batch effect correction methods have been developed. Linear model-based methods, such as limma (4), ComBat (5), and scMerge2 (6), gene-wisely estimate batch effects by including batch information as covariates in linear regression, and subsequently subtracting the estimates from gene expression matrix. While these methods can provide a full gene expression matrix after correcting for batch effects, they cannot consider cell type-specific effects in the correction process unless that information is explicitly provided. In contrast, similarity-based methods, such as Seurat-canonical correlation analysis (CCA) and reciprocal PCA (RPCA) (7), fastMNN (8), and Harmony (9), first embed cells in a shared low-dimensional space and match phenotypically similar cells across batches. They then compute local correction vectors to align these matched neighborhoods while preserving cell-type structure. However, this reliance on dimensionality reduction can lead to information loss and may restrict downstream analyses, while the computational methods also often incur high computational costs, limiting their scalability to large datasets. Generative model-based methods, such as scVI (10), produce a batch-corrected gene expression matrix by leveraging neural network architectures capable of capturing complex relationships among variables. Nevertheless, their ‘black box’ nature raises concerns about potential distortion of biological signals and the risk of overfitting cannot be ruled out. In addition to these limitations, to the best of our knowledge, existing batch-correcting algorithms do not explicitly address the potential distortion of variables of interest, such as disease states, which may occur when correcting for nuisance variables.

In this study, we introduce the Cell-type-specific Orthogonal Batch effect Removal Algorithm (COBRA), a method designed to remove batch effects from scRNA-seq data while preserving biological signals. It estimates a corrected full gene expression matrix, thereby ensuring the integrity of downstream analyses. COBRA offers several advantages: it (1) efficiently corrects batch effects based on a linear model, enabling its application to the large-scale scRNA-seq datasets; (2) accounts for heterogeneous batch effects across cell types by including interaction terms between batch and cell type variables in the model; and (3) preserves true biological signals by orthogonalizing batch-related variables with respect to the variables of interest. When cell type information is unavailable, COBRA estimates pseudo-cell types and incorporates these into the model. We validated the proposed algorithm extensively across a wide range of scRNA-seq datasets, including simulated datasets, benchmark datasets, and real-world studies, demonstrating the superiority of COBRA over existing methods.

## Results

### COBRA: Cell-type-specific, Orthogonal Batch effect Removal Algorithm

COBRA is designed to remove cell-type specific batch effects from scRNA-seq data using a linear model while preserving biological signals. The algorithm begins by constructing a design matrix that incorporates batch, cell type, the interaction between batch and cell type, and other relevant covariates such as disease status. COBRA then decomposes this matrix into batch-related and biological components. To ensure biological signals remain intact, the batch-related components are transformed to be orthogonal to the biological ones. These resulting orthogonalized batch variable are then used within a standard linear regression framework to estimate batch effects. Subtracting these estimated batch effects from the original expression matrix yields the final, batch-corrected expression matrix (**Figure 1a**, see **Methods**).

**Figure 1.**
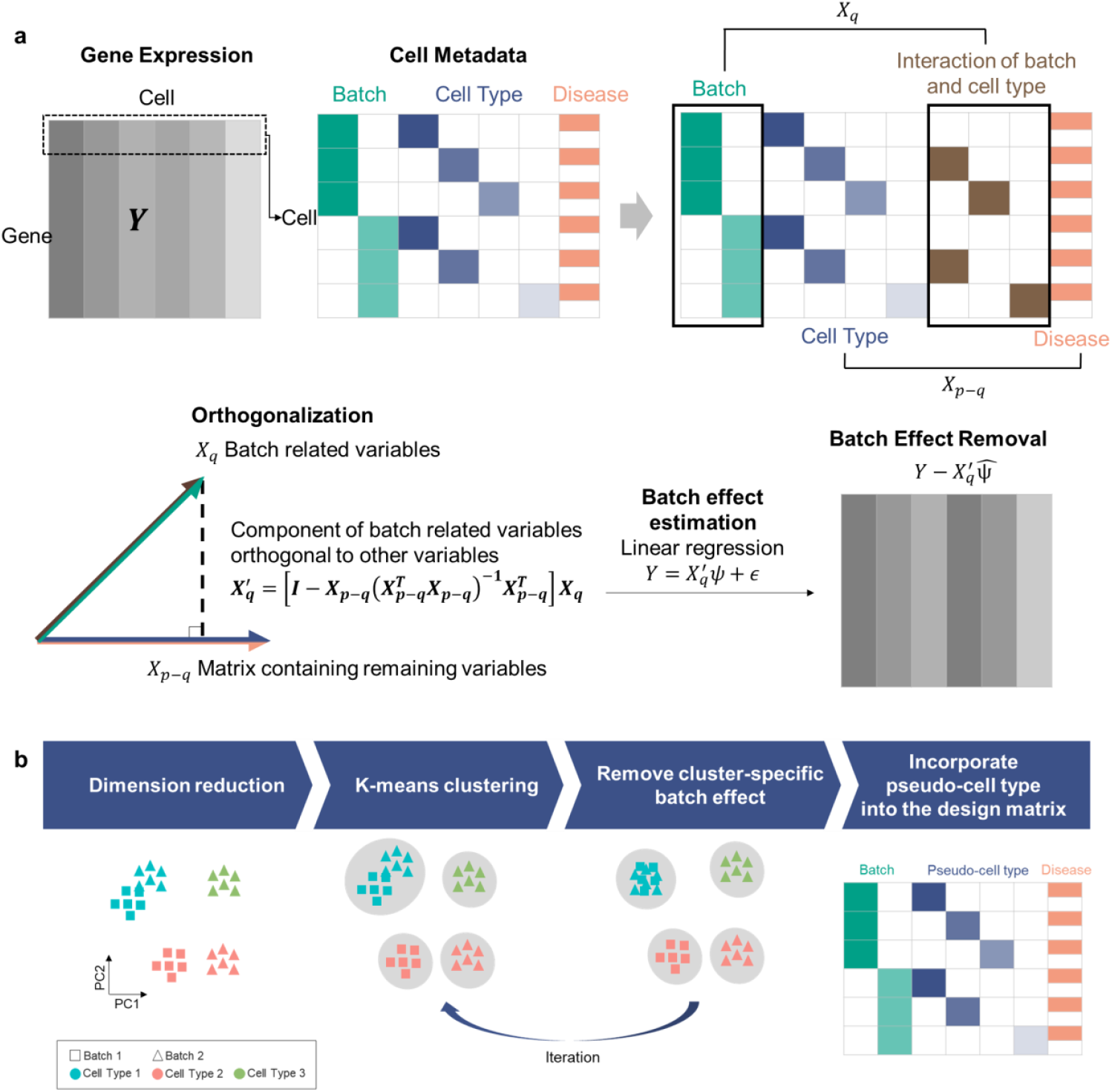
Overview of the COBRA algorithm. (a) Conceptual overview of the COBRA framework. A design matrix is constructed to include batch, cell type, interaction terms, and other covariates such as disease status. COBRA orthogonalizes batch-related variables with respect to biological signals. Batch effects are estimated using ordinary least squares (OLS), and the corrected gene expression matrix is obtained by subtracting the estimated batch effects from the original data matrix. (b) Workflow of pseudo-cell type estimation applied when cell-type labels are not available. The procedure iteratively clusters cells using K-means algorithm removes cluster-specific batch effects on dimension reduced matrix. The estimated clusters, referred to as pseudo-cell-types, are then incorporated into the design matrix in COBRA algorithm.

The output is a batch-corrected gene expression matrix with the same dimensions as the input data. Unlike embedding-based methods, which often reduce dimensionality and lose gene-level information, COBRA preserves the full gene expression matrix. This feature enables direct downstream analyses, such as differentially expressed genes and trajectory inference, without requiring additional processing steps or sacrificing analytical resolution.

In cases where cell type information is unavailable, COBRA implements an iterative clustering approach to estimate pseudo-cell types while accounting for batch effects. This procedure iteratively applies k-means clustering to dimension-reduced expression matrix to refine cluster assignments. In each iteration, linear regression is used to remove marginal batch effect across clusters, thereby optimizing the identification of cell populations in the absence of prior annotations. Subsequently, these estimated pseudo-cell type information are incorporated into the design matrix in place of known cell-type information. (**Figure 1b**, see **Methods**).

In summary, COBRA addresses the shortcomings of existing methods while maintaining practical utility for large-scale single-cell studies. By combining (1) cell type-specific batch modeling, (2) robust preservation of biological signals through orthogonalization, and (3) direct compatibility with downstream analyses, our approach provides a comprehensive solution for mitigating technical variation in scRNA-seq data without compromising biological insights. We applied COBRA and other existing batch correction methods to various benchmark datasets (see datasets section in **Methods and Materials**), and the key results are descripted in the subsequent sections, with additional findings included in **Supplementary Figure S5-10**.

### Cell-type-specific batch effect correction

We assume that batch effects in scRNA-seq data are not identical across cell types. To investigate this heterogeneity, we compared the average log2FC of gene expression between batches for each cell type. This analysis revealed that the average log2FC values differ significantly across cell types, with varying degrees of divergence depending on the experimental conditions (**Figure 2a**). This finding underscores the need for batch correction methods capable of addressing cell-type-specific variations to ensure reliable interpretation of scRNA-seq data.

**Figure 2.**
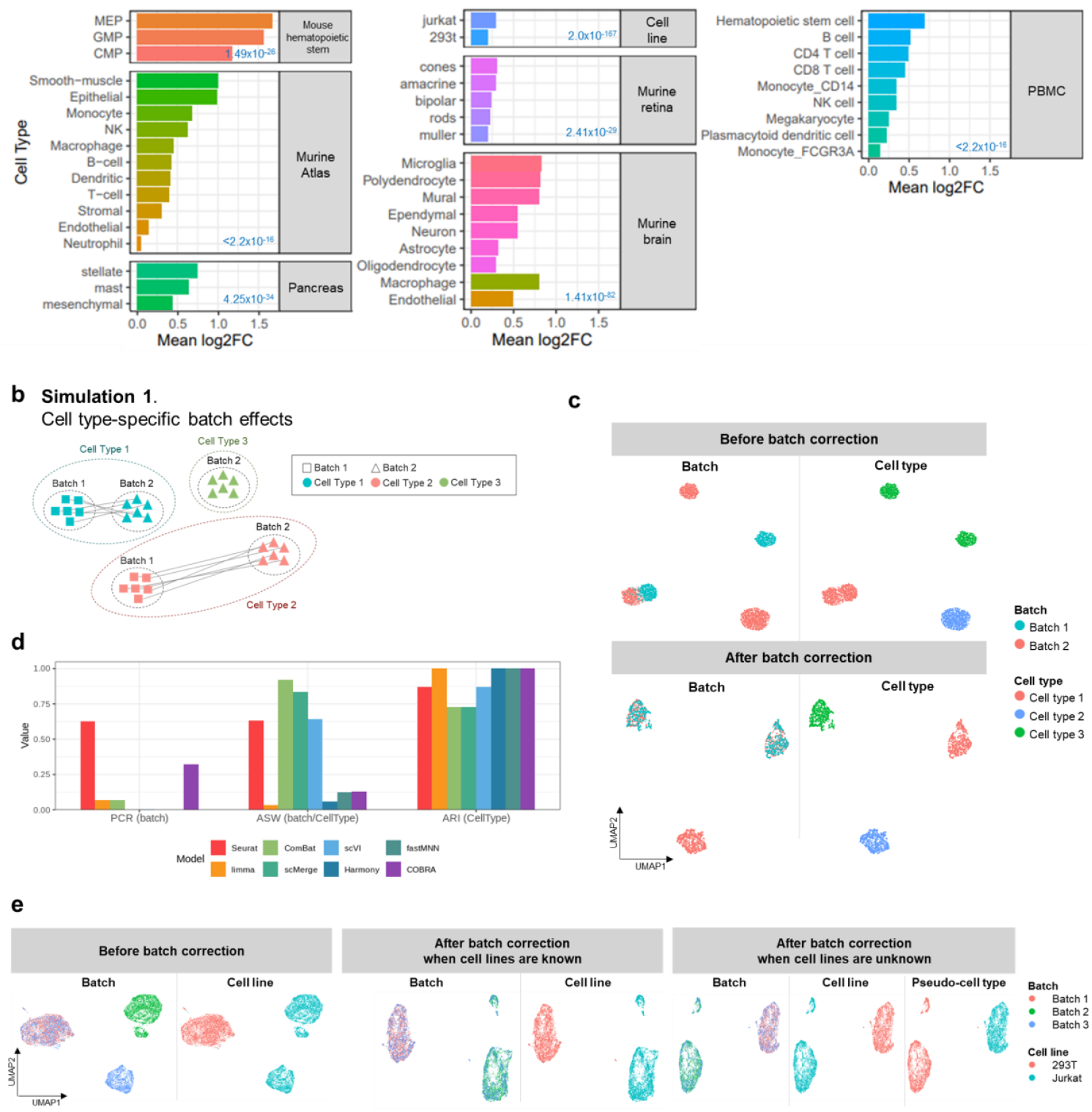
Cell-type-specific batch effects. (a) Variability of batch effect across scRNA-seq datasets. Batch effects are variable depending on cell type and dataset. Each bar indicates the average log2FC between two different batches within each cell type across scRNA-seq benchmark datasets. P-values from Kruskal-Wallis test are shown for each dataset. (b) Simulation scenario which has different batch effects across cell types. Cell type 1 show minimal batch effect, cell type 2 exhibits large batch effect, and cell type 3 appears in only one batch. (c) UMAP plots before and after batch correction using COBRA, colored by batch (left) and cell type (right). Post-correction results show batch integration while preserving biological clustering. (d) Quantitative comparison of batch correction methods using principal component regression (PCR) with R=50, average silhouette width (ASW), and adjusted Rand index (ARI). COBRA achieves a favorable balance between removing batch effects and maintaining cell type, compared to existing methods including Seurat, limma, ComBat, scMerge, scVI, Harmony, and fastMNN. (e)UMAP plots of a mixed cell line dataset before and after batch correction using COBRA. Corrections were applied both when cell line labels were available (middle) and unavailable (right). COBRA successfully integrates batches while retaining biological distinctions regardless of label availability.

To rigorously evaluate COBRA’s performance in correcting cell-type-specific batch effects, we simulated a scRNA-seq dataset comprising three distinct cell types: (1) a cell type with minimal batch effects, (2) a cell type with substantial batch effects, and (3) a cell type restricted to a single batch (simulation 1; **Figure 2b**). COBRA successfully integrated the batches while preserving the distinct biological structures of each cell type. This was evident in the uniform manifold approximation and projection (UMAP) visualization (**Figure 2c**), which demonstrated complete batch integration within individual cell types. We further benchmarked COBRA against widely used methods, including Seurat, limma, ComBat, scMerge, scVI, Harmony, and fastMNN. Performance was evaluated using both visual inspection of UMAP plots (**Supplementary Figure S1a**) and quantitative evaluation metrics (**Figure 2d)** (see evaluation metrics in **Methods and Materials**). When cell type labels were not provided, COBRA internally estimated pseudo-cell-type that perfectly matched the true cell-type (**Supplementary Figure S1a**), leading to identical downstream batch correction performance. These results demonstrate that COBRA effectively mitigates batch variability while preserving biological signals, outperforming existing approaches.

We further validated COBRA using a real dataset derived from two cell lines, Jurkat and 293T, distributed across three batches: batch 1 (293T only), batch 2 (Jurkat only), and batch 3 (a 50/50 mixture of both cell lines). In the uncorrected data, the 293T cells showed no clear separation by batch, whereas Jurkat cell displayed distinct clustering by batch **(Figure 2e**). The goal was to merge the Jurkat cells from batch 2 and 3 while maintaining the distinct cluster of 293T cells. COBRA successfully achieved this objective, outperforming the other methods (**Figure 2e** and **Supplementary Figure S1b**). Notably, when run without cell line information, COBRA’s internally estimated pseudo-cell types accurately corresponded to the true cell lines, resulting in an equally effective correction.

### Effective distinction of true biological signals from unwanted variations

We evaluated how well COBRA distinguishes true biological signals from unwanted technical variations by benchmarking it on simulated scRNA-seq datasets. These datasets were designed to reflect real-world experiments where batch effects are confounded with biological effects. Specifically, we varied the ratio of batch effect size to biological effect size and the proportion (*p*) of all cases assigned to the first batch (simulation 2; **Figure 3a**). A value of *p*=0 represents a completely confounded design, where one batch contains all controls and another contains all cases. Conversely, *p* = 0.5 represents a perfectly balanced design, with equal proportions of cases and controls within each batch.

**Figure 3.**
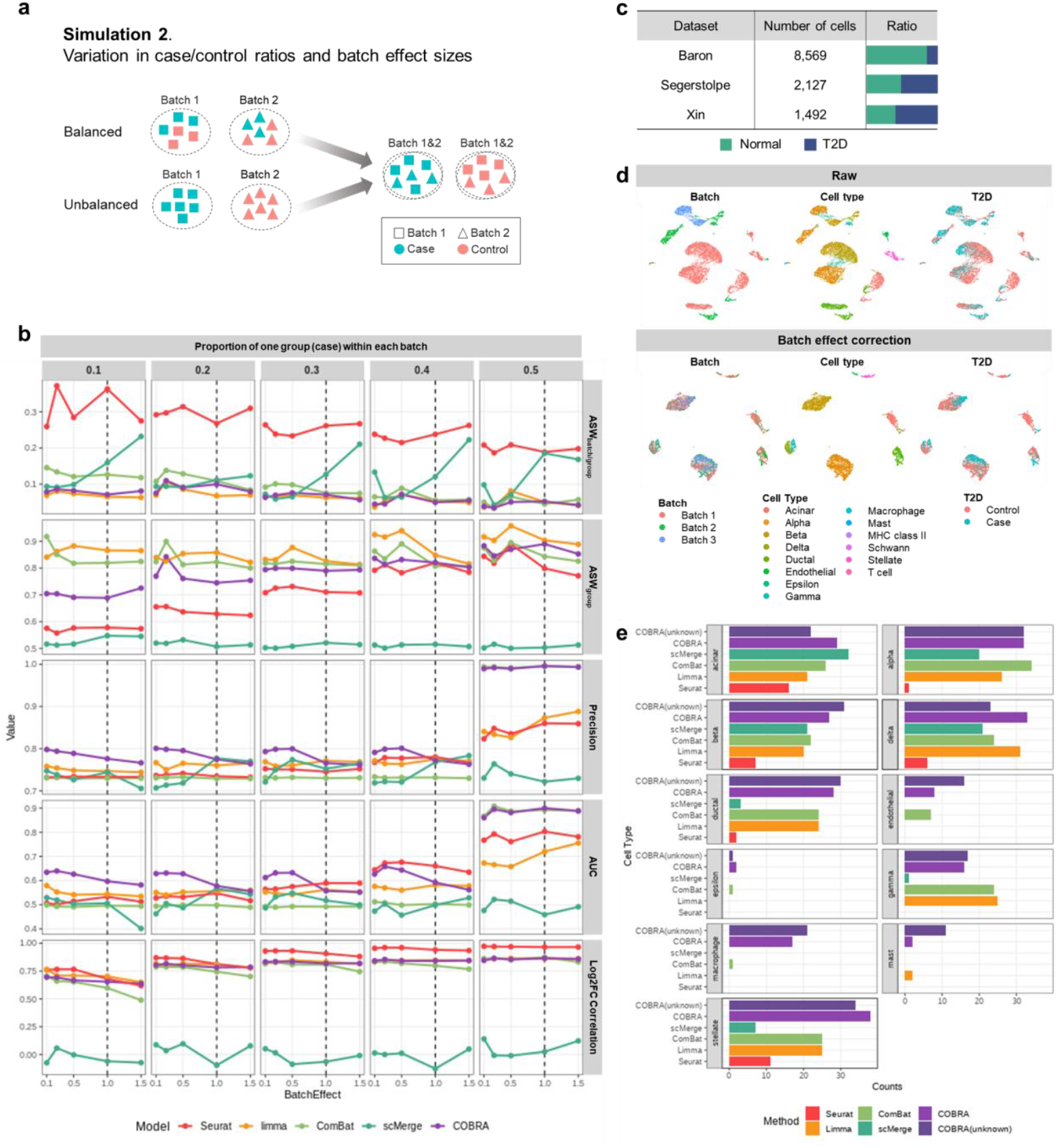
Distinction of true biological differences from unwanted variation. (a) Simulation scenario mimicking varying proportions of biological groups (case/control) within each batch. This setting models the situations where batch and disease status are partially confounded. (b) Comparative performance of batch correction methods under increasing the proportion of case within each batch. Evaluation metric include silhouette width based on batch within each group (ASWbatch/group), AWS for biological group separation (ASWgroup), and differential expression analysis accuracy: precision, AUC, and correlation of log2 fold changes for differentially expressed genes (DEGs) between before and after batch correction. (c) The number of cells and the proportions of cases and controls in each T2D dataset. (d) UMAP plot of T2D dataset before (top) and after (bottom) batch correction using COBRA. Color encodes batch (left), cell type (middle), and disease status (right). COBRA preserves biological separation while minimizing batch-driven clustering. (e)Total number of overlapped genes between DisGeNET (ID : C0011849) and detected DEGs across cell types. Comparisons are made between Seurat, limma, ComBat, scMerge, and COBRA, both with and without prior cell-type labels. COBRA consistently recovers a higher number of DEGs.

We assessed performance using evaluation criteria: (1) batch mixing efficiency within biological groups (Average Silhouette Width; ASW_batch/group_), (2) preservation of biological group structure (ASW_group_), and (3) three metrics related to differentially expressed gene (DEG) detection - precision, area under the curve (AUC), and the correlation between ground-truth and estimated effect sizes. The definitions of these metrics are provided in **Methods and Materials**. Ideally, batch-corrected data should exhibit clear separation by biological groups but strong mixing by batches while also enabling precise DEGs identification (high precision, AUC, and correlation values).

COBRA well separated cells by biological groups while effectively mixing cells from different batches across all conditions, as shown by its low ASW_batch/group_ and high ASW_group_ values (**Figure 3b** and **Supplementary Figure S2**). This indicates robust preservation of the biological group structure after batch correction. Notably, COBRA achieved superior performance in DEG detection, with the highest precision and AUC in nearly all scenarios, regardless of group proportions or relative effect sizes. The correlation between estimated and ground-truth effect sizes was also consistently high. Although the linear models without orthogonalization, such as limma and ComBat, exhibited relatively higher ASW_group_ values, their DEG detection metrics (precision and AUC) were lower than COBRA’s, indicating that excessive separation of groups may distort biological signals.

To further validate COBRA’s performance on biological signal prevention, we applied it to three T2D datasets with varying T2D-to-healthy subject ratios: Baron (0.85:0.15), Segerstolpe (0.49:0.51) and Xin (0.41:0.59) (**Figure 3c**). These datasets were treated as batches 1, 2, and 3, respectively, and were combined into a single integrated T2D dataset. After correction by COBRA, the UMAP plots showed a clear separation of cells by their phenotypic differences rather than by batch (**Figure 3d** and **Supplementary Figure S3**). To quantify the retention of biological effects, we performed DEG analysis by cell type, followed by gene set over-representation analysis using the DisGeNET (11), a curated database of disease-associated genes, as a source for T2D-related gene sets. COBRA identified a higher number of DEGs than existing methods did across most cell types, including those where other methods failed to detect any DEGs (**Figure 3e**). This higher sensitivity in detecting DEGs in both simulated and real datasets demonstrate COBRA’s ability to enhance biological discovery in scRNA-seq studies.

### Detection of differential cell states for multi-condition data

To assess COBRA’s capability to handle multi-condition and multi-batch datasets, we applied it to a large-scale COVID-19 dataset. Prior to batch correction, cells were clearly separated by batch. After applying COBRA, these batch-specific clusters were integrated, while maintaining cell type identities. Moreover, this correction enabled the identification of severity-associated cell states, such as B cells within specific immune populations. This suggests that these subsets may serve as sensitive indicators of disease progression (**Figure 4a** and **Supplementary Figure S4**).

**Figure 4.**
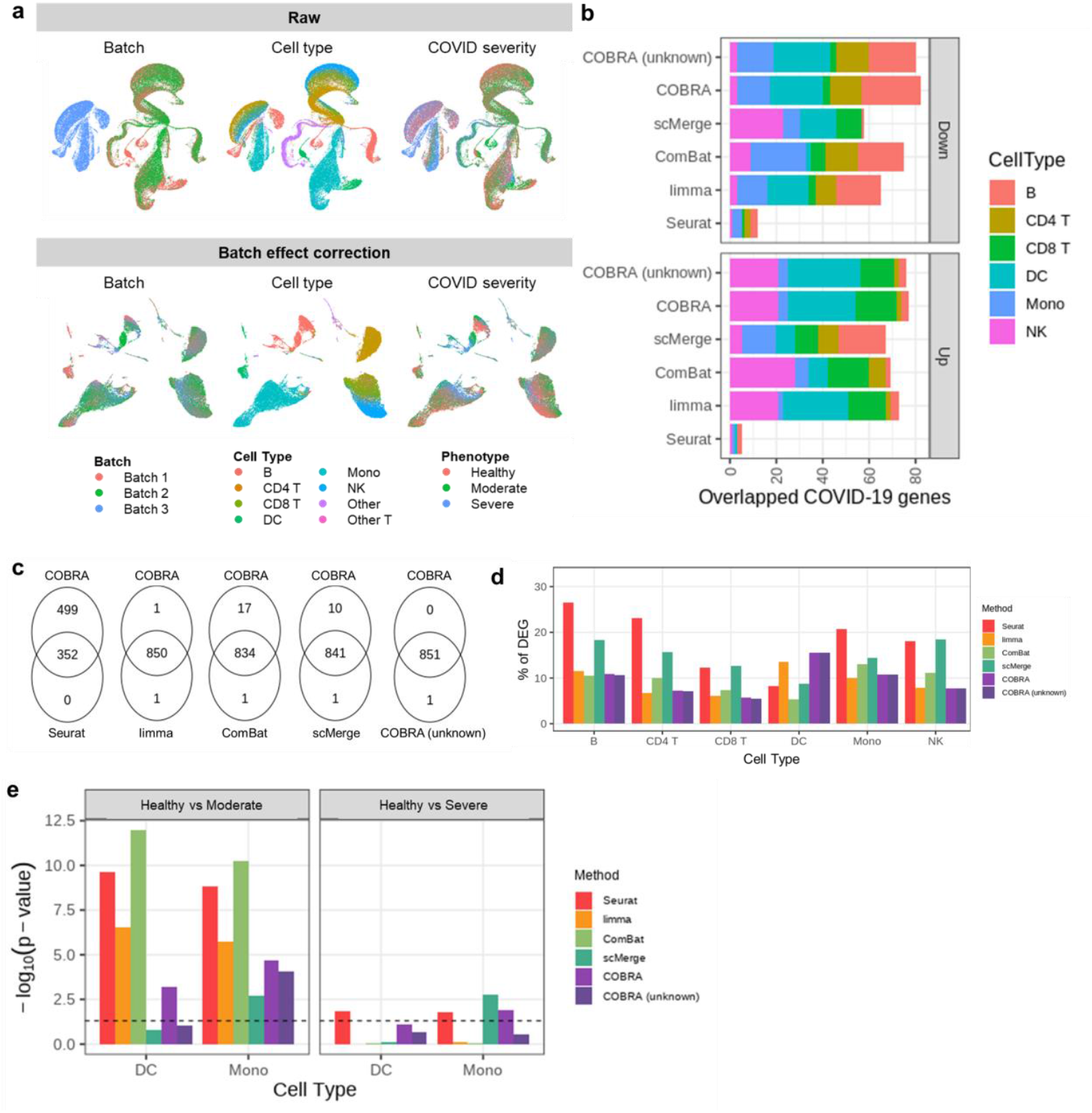
Identification of cell-state differences in COVID-19 across multiple severity groups. (a) UMAP plot of a COVID-19 peripheral blood mononuclear cell (PBMC) single-cell RNA-seq dataset before (top) and after (bottom) batch correction using COBRA. Cells are colored by batch (left), cell type (middle), and COVID-19 severity group (right). COBRA integrates batches while preserving biologically relevant structures. (b) Recovery of known COVID-19–related genes across methods. Bars represent the number of differentially expressed genes (DEGs) overlapping with previously published COVID-19 gene list. COBRA shows enhanced overlap for both down-regulated (up) and up-regulated (bottom) gene sets compared to other batch correction methods. (c) Venn diagrams comparing DEGs identified COBRA and other existing batch correction methods. (d) Proportion of DEGs among tested genes across cell types for each method. (e) Functional enrichment analysis of interferon (IFN) response genes between COVID-19 severity groups in dendritic cells and monocytes. The −log10(p-vlue) from a Kolmogorov–Smirnov test for Gene Ontology term “Response to type I interferon (GO:0034340)” is shown.

Subsequent analyses compared cell-type-specific DEGs with the previously reported bulk RNA-seq derived COVID-19 gene signatures (12). COBRA identified a greater overlap with known COVID-19 DEGs than existing batch correction methods, revealing which immune cell subsets exhibited predominant up- or down-regulation in response to infection (**Figure 4b**). COBRA recovered the DEGs detected by alternative methods, while also capturing additional COVID-19 associated genes that were not identified in elsewhere (**Figure 4c**). When quantifying the proportion of the detected DEGs, COBRA did not inflate the fraction of significant genes relative to existing approaches (**Figure 4d**), which may indicate that COBRA’s enhanced overlap is not driven by false positives, but rather reflects biologically relevant signals.

Further analysis focused on interferon (IFN) response genes (GO:0034340), which are known to be significantly altered in monocytes and dendritic cells following after the COVID-19 infection (13). COBRA’s results exhibited a significant enrichment with these IFN response genes across disease severity groups - healthy, moderate, and severe. While other algorithms showed significance in the ‘healthy vs. moderate’ comparisons only, COBRA additionally detected shifts between healthy and severe patients, consistent with findings from Edhiro et al. (14) (**Figure 4e**).

### Scalability of data integration

The linear model framework of COBRA ensures computational efficiency, with a complexity of *O*(*GTp*^2^), where *G* is the number of genes, *T* is the number of cells, and *p* is the number of coefficients in the design matrix. This structure enables efficient large-scale batch correction, providing advantage over more computationally demanding approaches such as similarity-based models with approximate complexity of *O*(*cT*^2^), where *c* is the number of reduced genes or dimensions and *c* ≪ *G*.

COBRA produce a full cells-by-genes matrix, ensuring direct compatibility with downstream analyses without dimensionality reduction. While this implies higher memory requirements, our benchmarking demonstrates that COBRA remains competitive in peak RAM usage when considering the entire pipeline, including preprocessing (**Figure 5a** and **5b**).

**Figure 5.**
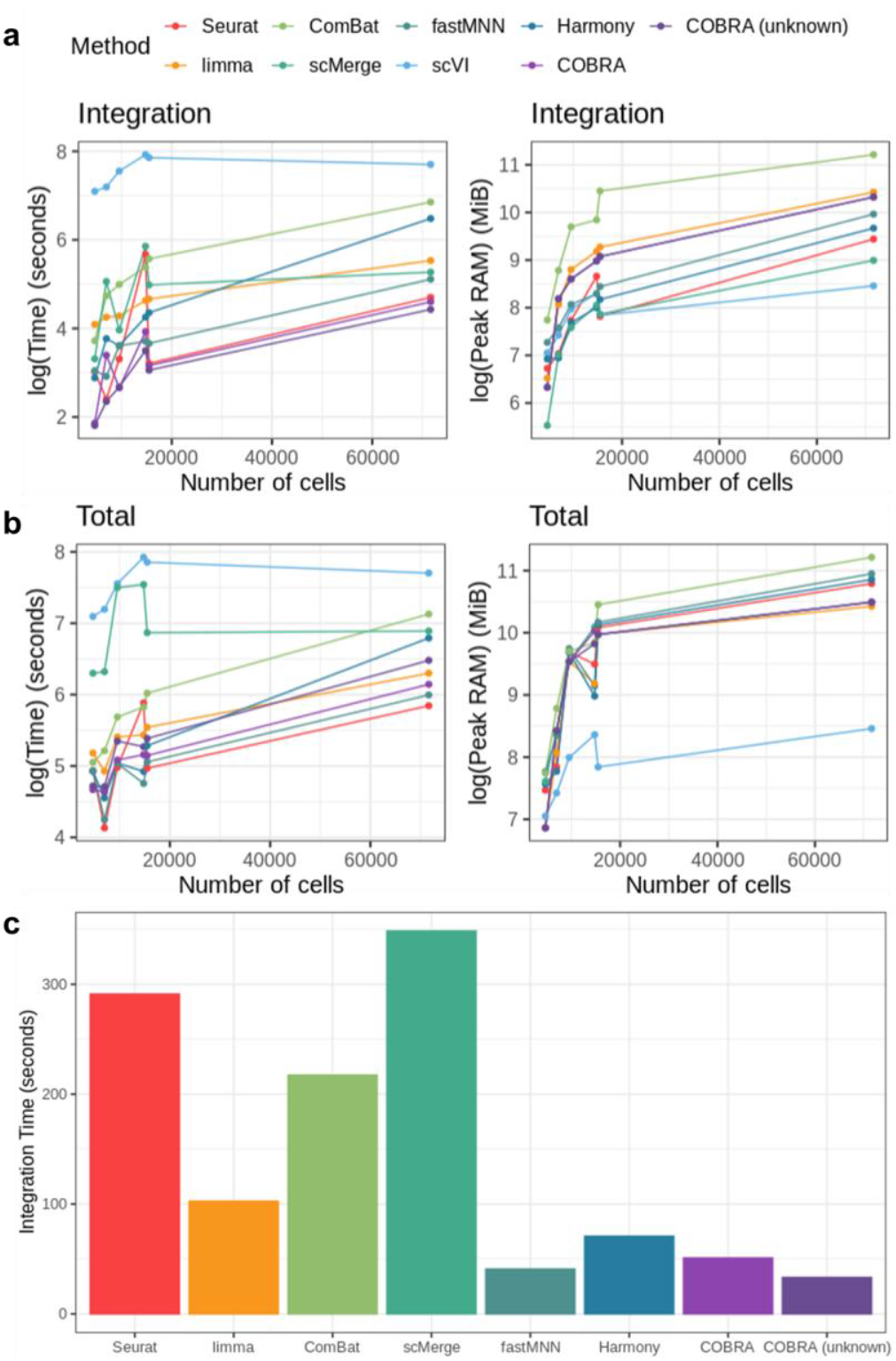
Computational scalability of COBRA and other batch correction methods. (a) Evaluation of integration runtime (left) and peak memory usage (right) across a range of dataset sizes (number of cells) during the integration step. COBRA demonstrate favorable scalability compared to existing methods including Seurat, limma, ComBat, scMerge, fastMNN, Harmony, and scVI. (b) Evaluation of total runtime (left) and peak memory usage (right) across a range of dataset sizes (number of cells) including both preprocessing and batch correction stages. (c) Integration runtime summary (in seconds) for each method for human pancreas dataset which have multiple batches.

Notably, COBRA showed the shortest integration time although it utilized the full matrix (**Figure 5a**). COBRA maintained competitive scalability, even when including preprocessing steps and pseudo-cell type estimation in the absence of prior cell type information (**Figure 5b**). Unlike methods that integrate batches sequentially, such as Seurat, COBRA provides a clear advantage when handling datasets with multiple batches (**Figure 5c**). Sequential integration methods require computational time that increases approximately with *T*^2^. In contrast, COBRA’s computational time increases only linearly with *T*, significantly improving runtime efficiency for large-scale data integration tasks.

## Discussion

scRNA-seq has revolutionized our ability to dissect cellular heterogeneity. However, batch effects remain a significant challenge to data integration and downstream analyses. To address this, we developed COBRA, a method that efficiently and effectively mitigates technical variation while preserving true biological signals. We validated our method using a variety of simulated and real scRNA-seq datasets, including those from studies of T2D and COVID-19. Across these diverse datasets, COBRA demonstrated robust performance in removing batch effects while maintaining cell-type identities and preserving biologically relevant gene expression pattern.

COBRA offers key advantages over widely used methods like Harmony and Seurat integration. While these methods rely on non-linear, embedding-based approaches, COBRA’s linear modeling framework provides a more interpretable and computationally efficient solution. This makes it particularly well-suited for the computational demands of single cell studies. COBRA is also able to effectively handle interest effects when correcting batch effects, which are often required in scRNA-seq data. Using a variety of simulated and real scRNA-seq datasets, including those from studies of T2D and COVID-19, we demonstrate that COBRA shows robust performance in removing batch effects while maintaining cell type separation and preserving biologically relevant gene expression patterns. These efficiency and interpretability make COBRA particularly appealing for large-scale studies where computational resources are a concern and clear, traceable results are paramount.

While COBRA demonstrates improvements, it shares fundamental limitations with all batch correction methods particularly when batch and phenotypic groups are completely confounded. This issue arises in real-world scenarios, such as when cases are processed in one batch and controls in another. In such cases, the biological signal of the disease and the technical batch effect are inseparable, making it impossible to distinguish between them. Even with COBRA’s orthogonalization approach, this inherent confounding can lead to the removal of true biological differences. Future research could explore advanced statistical models or Bayesian frameworks that might better handle such scenarios by incorporating a priori knowledge or prior distribution about the expected biological variance.

A second limitation of COBRA is its reliance on K-means clustering for pseudo-cell type estimation when cell type information is not available a priori. Although K-means clustering is computationally efficient, its effectiveness depends on well-separated clusters in the data (15). This can pose challenges when cell types exhibit overlapping transcriptional profiles or when other biological effects dominate the clustering structure (16). Specifically, when strong phenotypic or environmental influences overshadow cell-type-specific signals, K-mean clustering may produce clusters based on those effects rather than true cell identities, potentially masking batch effects within clusters. Incorporating prior knowledge about cell type identification using marker genes or mapping algorithms to publicly available single-cell atlases (17) could improve accuracy of clustering and downstream correction.

## Materials and methods

### COBRA

#### Batch effect estimation

COBRA incorporates batch information, cell types, and any covariates into the design matrix to systematically account for batch-related variations. To model cell-type-specific batch effects, we introduce an interaction term between batch and cell type in the design matrix. If cell type annotations are unavailable, COBRA can estimate pseudo-cell types using iterative clustering algorithm described in the **Figure 1b**. COBRA divided the design matrix into batch-related and batch-unrelated matrices, and then batch-related variables are orthogonalized with respect to other variables to preserve biological variability. Subsequently, scRNA-seq expression data is regressed on orthogonalized batch-related variables, and the resulting residual expression is used as the batch-corrected data.

Prior to batch correction, COBRA requires normalization and scaling for unique molecular identifier (UMI) count by batch. Let *G* be the number of genes and *B* the number of batches. In batch *b*, subject *s* has *τ*_*bs*_ cells, giving the total number of cells 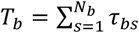 across *N*_*b*_ subjects. Overall, across batches, the total number of subjects is 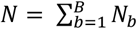 and the total number of cells is 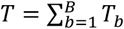. For gene *g* in cell *j* of subject *s* sequenced in batch *b*, let the raw UMI count be 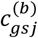 and the cell’s library size be 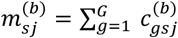. Within each batch, counts are normalized as

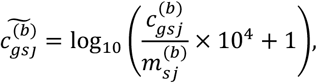

and then scaled per gene via

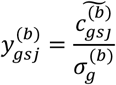

where 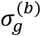 is the standard deviation of 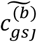 for gene *g* in batch *b*.

Then, for gene *g*, COBRA fits

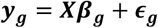

where 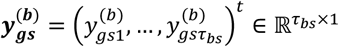 denotes the cell-level vector for subject *s* in batch 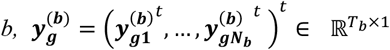 includes all subjects within batch *b*, and 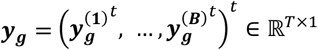 has expression vectors across batches. The design matrix ***X*** includes batch, cell type, batch-cell type interaction terms, and phenotypic covariates, having *p* variables so ***X*** ∈ ℝ^*T*×*p*^. The corresponding coefficient vector is expressed as ***β***_***g***_ ∈ ℝ^*p*×1^, and ***𝝐***_***g***_∼N(0, σ^2^***I***_**T**_) ∈ ℝ^*T*×1^ represents the random error, where *I*_*T*_ denotes the *T* × *T* identity matrix.

By partitioning the ***X*** into 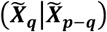, where 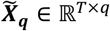 is a matrix containing the batch-related variables, such as batch and the interaction variables, and 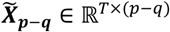 is a matrix containing the other variables, we can write ***Xβ***_***g***_ as 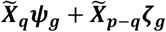, as following (18)’s notation. If we transform 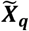 into 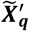, where 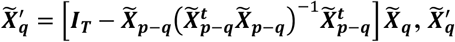 becomes orthogonal to 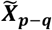 and we have 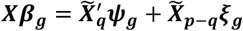 where 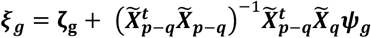. Consequently, the coefficients ***𝝍***_***g***_ and ***ξ***_***g***_ can be estimated using least square method, resulting in the expression of gene *g* with the batch effect removed: 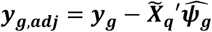 (**Figure 1**).

COBRA’s computational complexity is determined by three key steps. First, for each gene, the orthogonalization of batch-related variables requires *O*(*T*(*p* − *q*)^2^ + (*p* − *q*)^3^ + *T*(*p* − *q*)*q* + (*p* − *q*)^2^*q*). The batch effect estimation and batch effect removal require *O*(*Tq*^2^) and *O*(*Tq*), respectively. Considering *T* ≫ (*p* − *q*) and *T* ≫ *q*, the total computational complexity for all the genes is approximately *O*(*GTp*^2^), showing the complexity only linearly increases as *G* or *T* increases.

#### Pseudo-cell type estimation

For datasets where cell type information is unavailable, COBRA employs an iterative approach to estimate pseudo-cell types. This approach simultaneously identifies biologically meaningful cell clusters and corrects for batch effects.

The top 20 principal component (PC) scores of each cell are calculated using selected featured genes across batches. The process begins by clustering a random subset (10%) of cells to enhance computational efficiency and reduce noise. K-means clustering is then performed on the PC scores, with the optimal number of clusters determined by maximizing silhouette score. The resulting cluster assignments are then used to classify the remaining cells into these initial groups. Next, these clusters are treated as preliminary pseudo-cell types, and the PC scores are adjusted for batch effects using the orthogonalization-based correction method described in the *Batch effect estimation* section. This initiates an iterative refinement cycle. In each iteration, the batch-corrected PC scores for all cells are re-clustered using K-means, again determining the optimal number of clusters by maximizing the silhouette score. These new clusters serve as updated pseudo-cell types, which are then used to re-estimate the pseudo-cell types and correct for batch effects on the PC scores. This iterative process is repeated until the model fit, as measured by Bayesian information criterion (BIC), no longer improves or until a maximum of 20 iterations is reached. Once a stopping criterion is met, the algorithm outputs the final pseudo-cell type assignments, ensuring that the resulting clustering structure is robust to batch artifacts.

### Other integration methods

To comprehensively evaluate COBRA against existing batch correction methods, we implemented standard integration pipelines for widely used scRNA-seq batch correction methods. Below, we describe the preprocessing and integration steps followed for each method.

#### Seurat

We followed the recommended integration pipeline from Seurat v5 (7). First, each batch was pre-processed individually by normalizing the data, identifying highly variable genes, and scaling gene expression values. Principal component analysis (PCA) was then performed across all batches. These dimension-reduced datasets were then used to identify anchors for integration. During anchor identification, we considered both CCA and RPCA for comparison. The output was a batch integrated dataset in a low-dimensional space, which was subsequently projected into UMAP space for visualization and assessment. Comparing the phenotypic effect between integration methods requires a full gene-by-cell expression matrix for downstream analyses like DEG testing. Therefore, for such comparisons, we employed full matrix for integration.

#### Harmony

The Harmony (9) integration pipeline followed the standard approach as typically implemented in Seurat. The initial steps involved normalizing data and identifying highly variable genes. Then, the scaled data underwent dimensionality reduction into the PC space. Integration was performed by applying Harmony to the PC embeddings using the *IntegrateLayers* function with the *method=HarmonyIntegration* option within the Seurat package. Similar to Seurat, the integrated output was a reduced-dimensional embedding, which was used for UMAP visualizations.

#### fastMNN

For fastMNN (8) integration, preprocessing steps were identical to those of Seurat, including normalization, finding high variable genes, scaling and PCA. Then, batch correction was performed using fastMNN via the *IntegrateLayers* function with *method=FastMNNIntegration*. fastMNN provides batch-corrected data in a reduced-dimensional space.

#### ComBat

Since ComBat (5) is a linear model-based batch correction method, we applied identical preprocessing steps as for COBRA, such as batch-wise normalization and scaling. Batch correction was performed using the *ComBat* function specifying batch variable in the *batch* option and including other covariates, such as cell type and phenotype, as part of the design matrix in the *mod* option.

#### limma

Like ComBat, limma (4) is a linear model-based method and preserves the full gene matrix. The *removeBatcheffect* function was used with the same preprocessing steps and design matrix as used in COBRA.

#### scMerge

The scMerge (6) pipeline begins with normalization and scaling across the entire dataset. Stably expressed genes (SEGs) were identified as negative control genes by selecting those with an *scSEGIndex90* score of less than 0.5. Using these genes along with batch and cell type information, we performed batch effect removal with the *scMerge2* function.

#### scVI

The scVI model (10) is based on a conditional variational autoencoder. It employs raw count data for negative binomial modeling, and we trained the model for a maximum of 20 epochs. From the trained model, a latent presentation for each cell was extracted. This output is a multi-dimensional embedding where the batch effects have been removed.

### Datasets

Both experimentally derived and simulated datasets were used to evaluate the performance of COBRA compared to other batch correction methods. The benchmark datasets were obtained from https://hub.docker.com/r/jinmiaochenlab/batch-effect-removal-benchmarking which were originally introduced in Tran et al. (19). Annotated cell types from these datasets were used directly in our study.

#### Benchmark datasets

##### Cell line

The dataset comprises three sets of data (20, 21), each with a specific cell type composition. The first dataset consists of 2,885 293T cells, the second dataset includes 3,258 Jurkat cells, and the third dataset consists of a 50:50 ratio of Jurkat to 293T cells, totaling 3,388 cells. A total of 32,738 genes were obtained using the 10x Genomics platform and included in the gene expression dataset. The dataset is utilized to evaluate batch correction on monoclonal populations of each physiologically distinct cell type.

##### Human pancreas and T2D

The human pancreas dataset was acquired from five distinct sources (22-26), consisting of pre-processed count data with a total of 14,767 cells. These cells belong to 15 cell types, some of which are not shared across all batches. This dataset provides insight into the integration of multiple batches containing non-identical cell types. Among the five datasets, three have phenotype information indicating whether the patients are healthy or have T2D. We used these three datasets for evaluating the ability to preserve biological effects. ***Murine atlas*** Two distinct mouse cell atlas datasets were independently generated by different research groups using different scRNA-seq technologies. The first dataset was produced by Han et al. (27) through Microwell-Seq, and the second by the Tabula Muris Consortium (28) using Smart-Seq2 protocols. We integrated a total of 6,945 cells representing common 11 cell types from diverse organ systems and 15,006 shared genes across both datasets. Moreover, samples were obtained from nine murine organ systems - urinary, digestive, respiratory, circulatory, muscular, immune, nervous, endocrine, and lymphatic – allowing for comparisons of multiple cell types across different tissues. This dataset is intended to evaluate the batch effect removal across varying tissues and different scRNA-seq technologies.

##### Human PBMC

Two distinct datasets were acquired from healthy donor peripheral blood mononuclear cells (PBMCs) using the 3’ and 5’ 10x Genomics protocols, which are designed to capture different regions of mRNA (21). The first dataset consists of 8,098 cells while the second contains 7,378 cells for identical cell types, and both include 17,430 genes. This dataset allows for an investigation into how well batch correction performs when there are biological variations driven by different protocols.

##### Mouse retina

These data were obtained using drop-seq technology from two independent groups (29, 30), comprising 26,830 cells in the first batch and 44,808 cells in the second batch. Each cell contains expression values for 12,333 genes. The first batch contains twelve cell types, while the second batch contains five, which are a subset of the cell types from the first batch. The dataset was used to test batch correction on a dataset with non-identical cell types. Additionally, the original data exhibited minimal batch separation, and we investigated whether these patterns remained after applying integration algorithms.

##### Mouse hematopoietic stem and progenitor cells

The dataset was obtained by Nestorowa et al. (31) and Paul et al. (32), which were sequenced using the SMART-seq2 and MARS-seq protocols, respectively. From the raw datasets, we retained only cells with curated cell-type labels provided by Tran et al. (19), resulting in 1,920 and 2,729 cells in each batch and 3,467 common genes. This dataset was used to evaluate how well batch correction techniques mitigate batch effects caused by variations in sequencing technology when the cell compositions are different.

##### Mouse brain

This dataset combines two mouse brain datasets from Saunders et al. (33) and Rosenberg et al. (34) to evaluate the effectiveness of batch correction strategies on a large-scale dataset comprising 833,206 cells, with 691,600 cells in the first batch and 141,606 in the second batch. The primary objective is to assess the scalability of batch correction methods on large datasets.

##### COVID-19

The FRED HUTCH (35) provides standardized scRNA-seq datasets from published studies to enhance the understanding of COVID-19. To test integration methods in situations involving multiple phenotypic effect, we obtained three datasets - ARUNACHALAM, LEE and WILK - each with available phenotypic information. The metadata includes disease status; healthy, moderate or severe COVID-19. The data comprises six healthy individuals and nine patients with COVID-19, seven of whom were classified as having a moderate condition and two as severe. The dataset contains 175,345 cells and 33,516 genes and was used to assess batch effect removal methods for handling multiple phenotypes.

#### Simulation datasets

We generated synthetic scRNA-seq data using the splatter R package (36) to assess performance on two specific challenges: cell-type-specific batch effect and confounding batch effect.

The first scenario was designed to investigate the differential impact of technical variations across cell populations. We generated three distinct cell populations, each comprising 1,000 cells and 1,000 genes with varying degrees of batch effects. The magnitude of batch effects was controlled through two key parameters: *batch*.*facLoc*, which determines the average shift in expression values between batches, and *batch*.*facScale*, which modulates the variability of these batch-specific changes. Cell type 1 was assigned minimal batch effects (*batch*.*facLoc* = 0.05, *batch*.*facScale* = 0.05), cell type 2 was assigned substantial batch effects (*batch*.*facLoc* = 0.5, *batch*.*facScale* = 0.5), while cell type 3 was restricted to a single batch, which indicates no batch effect. After applying batch correction methods, principal component regression (PCR) and ASW were calculated to measure the degree of cell-type separation and the extent of batch mixing within the corrected datasets.

The second scenario addresses phenotypic imbalance across experimental conditions. Using parameters derived from the Camp reference dataset (37), we simulated 5,000 cells across 1,000 genes, distributed equally between two batches. To assess performance under these imbalanced conditions, the proportion of case cells within the first batch was varied from 0.1 to 0.5. Differential expression between case and control conditions was implemented uniformly across batches, *de*.*facLoc* = 1 and *de*.*facScale* = 1, where these parameters control the location and scale of the differential expression factors, respectively. The relative batch effect sizes were modulated by *batch*.*facLoc* parameters, which was varied from 0.1 to 1.5. The performance of batch correction methods was evaluated using ASW, precision, AUC and correlation of log_2_FCs for DEGs between before and after batch correction.

### Evaluation metrics

#### ASW

ASW was used to measure the quality of clustering in terms of both compactness and separation. It evaluates how similar an object is to its own cluster compared to other clusters by calculating the average silhouette width of all cells in the data. The silhouette score, *s*(*j*), for an individual cell *j* and the overall ASW for a dataset of *T* cells are defined as:

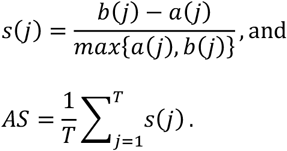

Here *a*(*j*) is the average distance from cell *j* to the other cells in the same cluster, and *b*(*j*) is the smallest average distance from cell *j* to cells in a different cluster. A higher ASW score suggests well-defined and distinct clusters, with a value approaching 1 indicating optimal clustering.

Additionally, we utilized ASW to assess the degree of separation among batches or biological groups within each cell type. Let *M* be the set of cell types, and for *m* ∈ *M* let *C*_*m*_be the set of cells belonging to the cell type *m*. Within each *C*_*m*_, *s*(*j*) is computed by treating batch labels as clusters. Following Luecken et al. (38), ASW is calculated as a weighted average across all cell types:

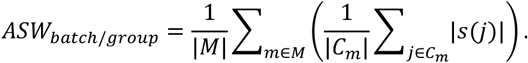

Here |·| denotes the number of elements in the set. ASW_batch/group_ ranges from 0 to 1, and a score close to 1 indicates clear separation, whereas a score close to 0 indicates poor separation.

#### PCR

To quantify the variance in the data attributable to batch effects, we utilized a score based on PCR. This score is calculated by summing the variance explained by top *R* PCs, where each PC’s variance is weighted by the proportion of its variance that is explained by the batch variable.

Let *z*_*r*_ denote the score vector for the *r*-th PC scores. The term *R*^2^(*z*_*r*_|batch) represents the coefficient of determination from a linear model regressing the batch variable on *z*_*r*_. Then, the total variance explained by the batch, *Var*(***y***_***g***_|*batch*), is then given by (38):

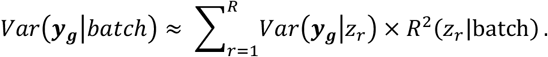

Here *Var*(***y***_***g***_|*z*_*r*_) is estimated from the mean squared regression of ***y***_***g***_ on *z*_*r*_. A higher *Var*(***y***_***g***_|*batch*) indicates a stronger batch effect, while a large reduction in this score after correction reflects more effective batch removal. All calculations of *Var*(***y***_***g***_|*batch*) were performed using the Python (21) module (38).

#### Precision, AUC and log_2_ FC

To evaluate the performance of each batch correction method on the simulation datasets, we assessed its ability to preserve true DEG signals. Performance was quantified using precision, AUC and log_2_ *FC*. Precision was defined as the proportion of correctly identified DEGs among all detected DEGs; a higher precision indicates a more effective distinction between true signals from false positives. We also computed AUC-ROC score to measure the overall ranking performance. Finally, to determine if the magnitude of the biological signal was maintained, we calculated the correlation of log_2_ *FC*s for true DEGs before and after batch correction.

#### Time and memory

We benchmarked the execution time and peak memory usage of our method in a single-core Linux environment. Execution time was measured using the *bench_time* function from the bench R package (39). Peak memory consumption was assessed using the *peakRAM* function from the peakRAM R package (40) with the maximum usage recorded from the {Peak_RAM_Used_MiB} output field.

### Analysis of DEGs

After applying batch correction, we performed differential gene expression analysis for each cell type individually. This analysis was conducted using *FindMarkers* function from the Seurat R package, with implementing the MAST algorithm (41). Genes were considered DEGs if they met the criteria of an adjusted p-value < 0.05 and |log2 fold change| > 1.5.

To assess whether biologically relevant signals were preserved after batch correction, the identified DEGs were compared against external disease-associated gene lists. Specifically, we used COVID-19 associated genes reported in Xiong et al. (12) and T2D associated genes obtained from DisGeNET (ID : C0011849) (11). The overlap between the detected DEGs and these disease-associated gene lists was used to evaluate the extent to which disease-relevant gene were obtained after batch correction.

### Gene-set enrichment analysis

The functional relevance of the identified DEGs were examined by gene-set enrichment analysis. The genes were ranked by differential expression p-values, and enrichment significance against on Gene Ontology (GO) term was evaluated using Kolmogorov-Smirnov (KS) test implemented in the topGO R package (42).

## Abbreviations

ASW: Average silhouette width
AUC: Area under the curve
BIC: Bayesian information criterion
CCA: Canonical correlation analysis
DEG: Differentially expressed gene
IFN: Interferon
KS: Kolmogorov-Smirnov
MAST: Model-based analysis of single-cell transcriptomics
PBMC: Peripheral blood mononuclear cell
PC: Principal component
PCA: Principal component analysis
PCR: Principal component regression
RPCA: Reciprocal principal component analysis
UMAP: Uniform manifold approximation and projection
UMI: Unique molecular identifier

## Declarations

### Availability of data and materials

The benchmark datasets used and/or analysed during the current study are available in Tran et al. (19), https://hub.docker.com/r/jinmiaochenlab/batch-effect-removal-benchmarking. The codes are publicly available at Zenodo (https://doi.org/10.5281/zenodo.17825327)

### Authors’ contributions

S.S, S.W and K.P contributed to study design. S.S and K.P participated in the analysis and interpretation of the results including benchmarking and simulation. S. S wrote the draft manuscript and S. W and K.P supervised the study and revised the manuscript. All authors have approved the final version of the manuscript for publication.

