## Supplementary for "COBRA: Cell-type-specific Orthogonal Batch effect Removal Algorithm in single cell RNA-sequencing data"

**Supplementary Figure S1. UMAP visualization of batch integration for (a) simulation 1 dataset and (b) cell line dataset**

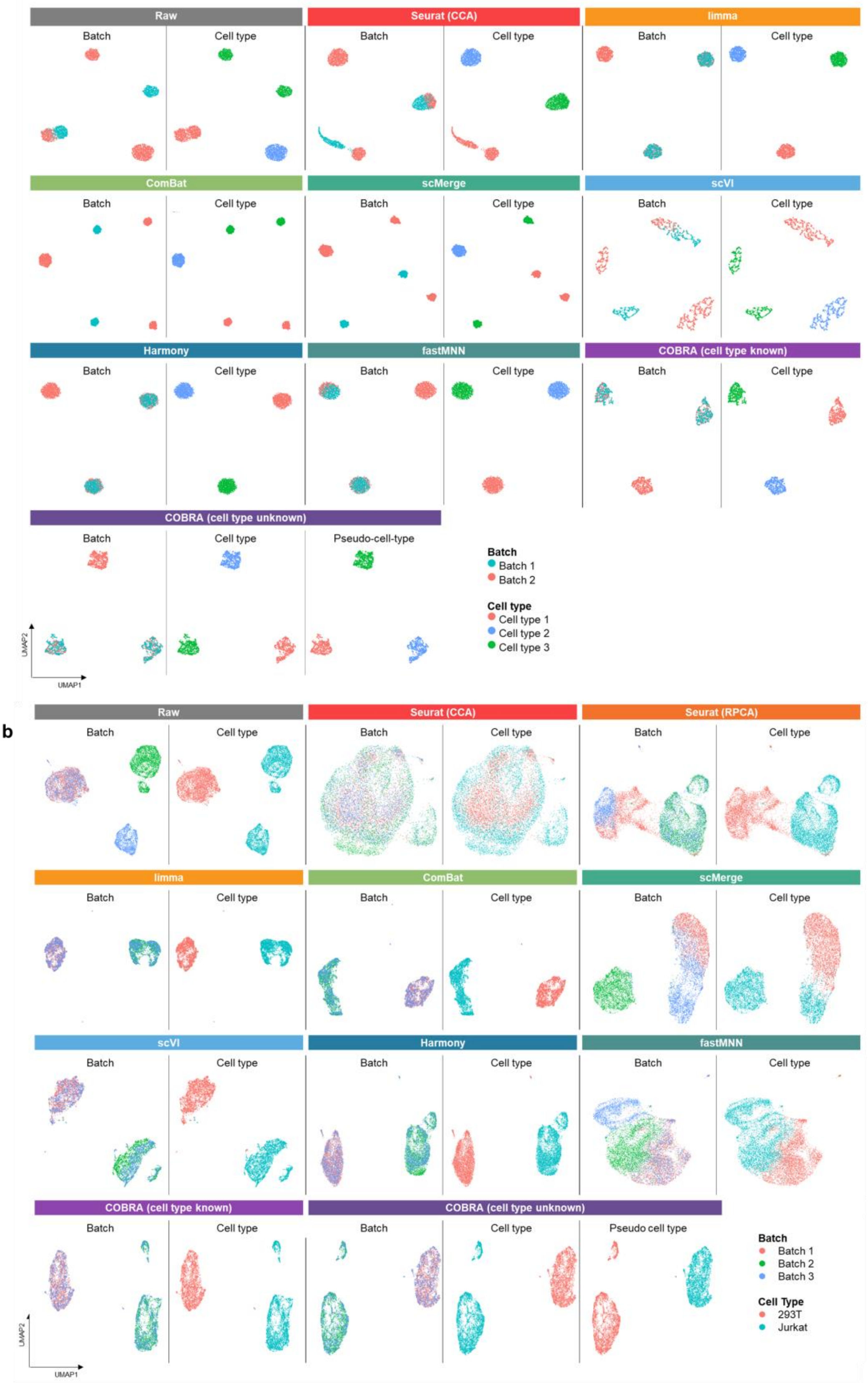

**Supplementary Figure S2. UMAP plot of batch integration for simulation 2 (a) raw dataset and (b) COBRA integrated dataset by batch (left) and cell type (right)**

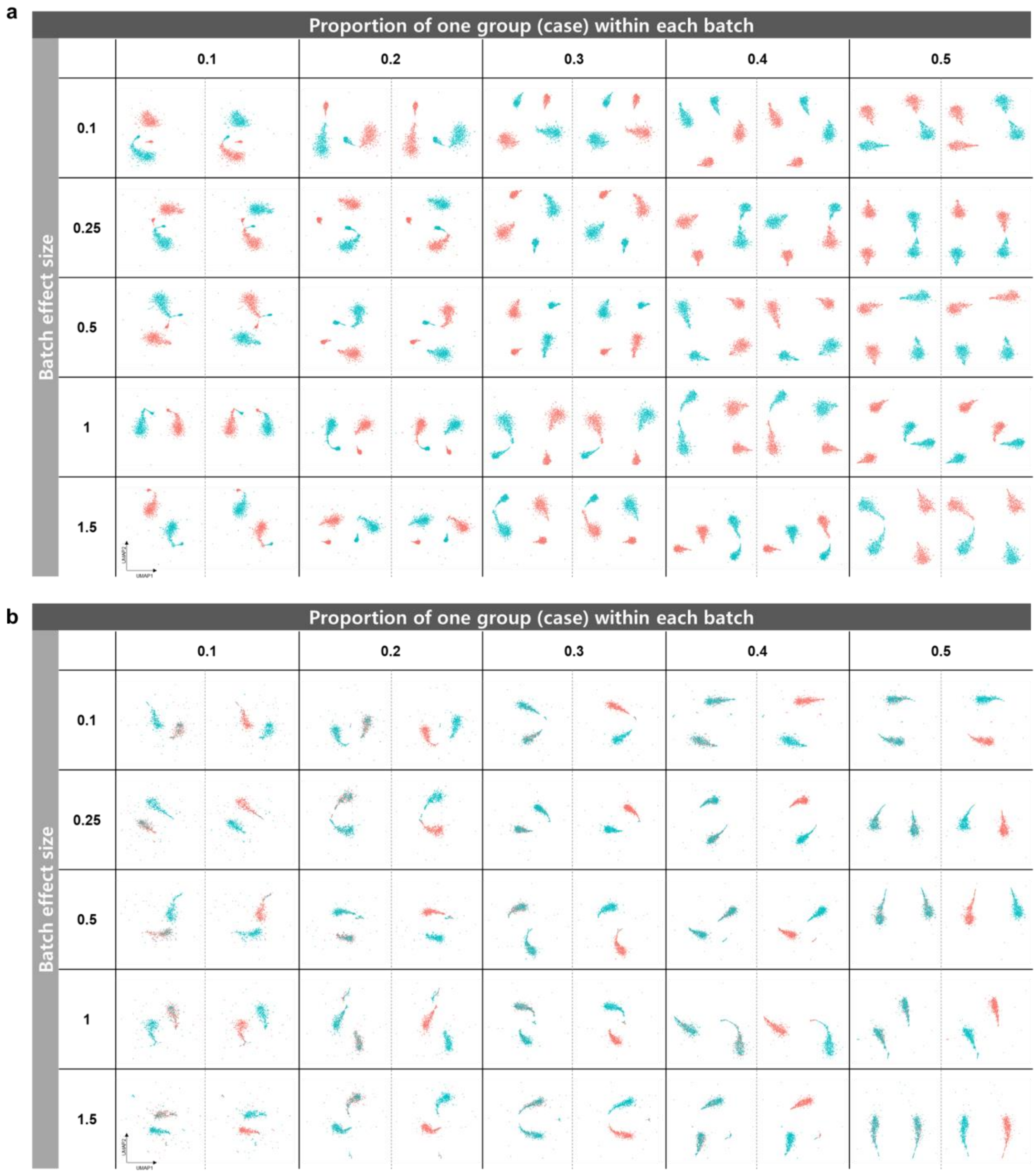

#### Supplementary Figure S3. UMAP visualization for T2D dataset

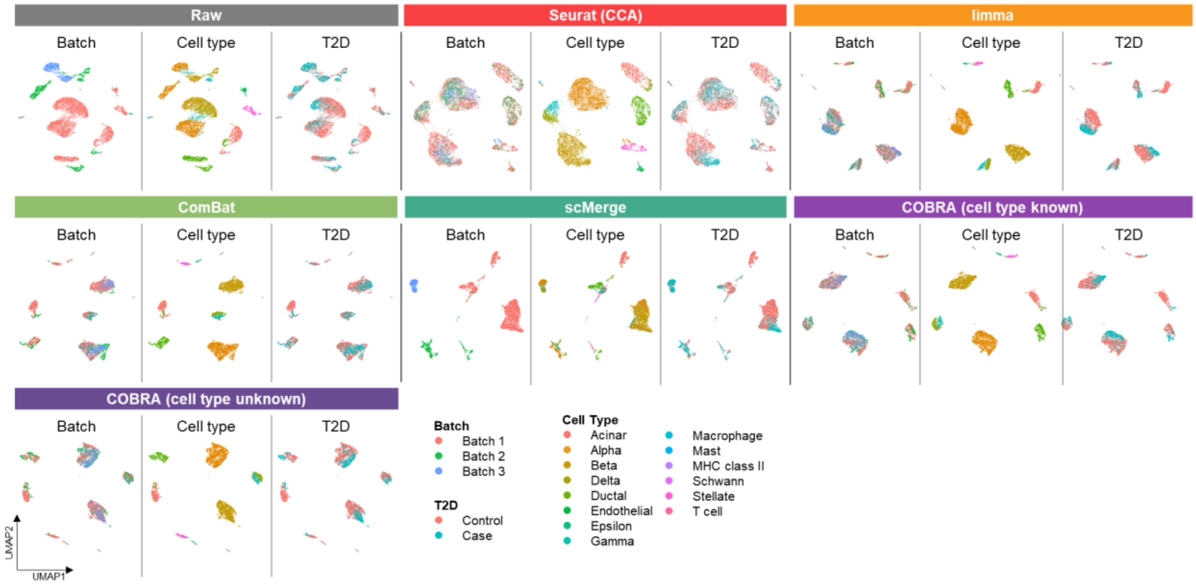

#### Supplementary Figure S4. UMAP visualization for COVID-19 dataset

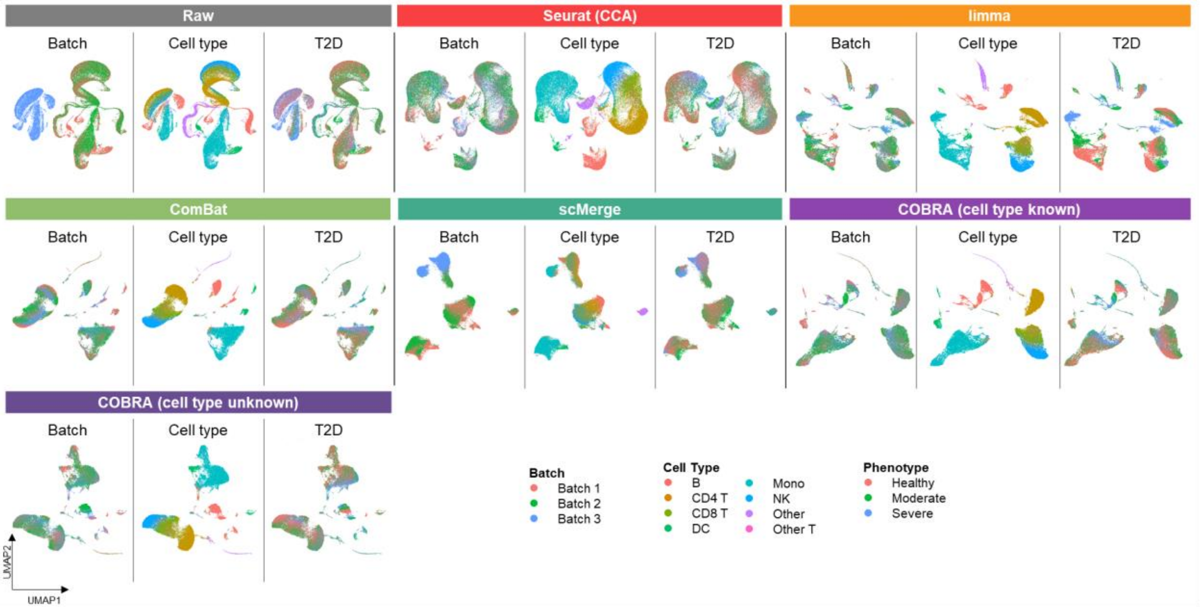

#### Supplementary Figure S5. UMAP visualization for murine atlas dataset

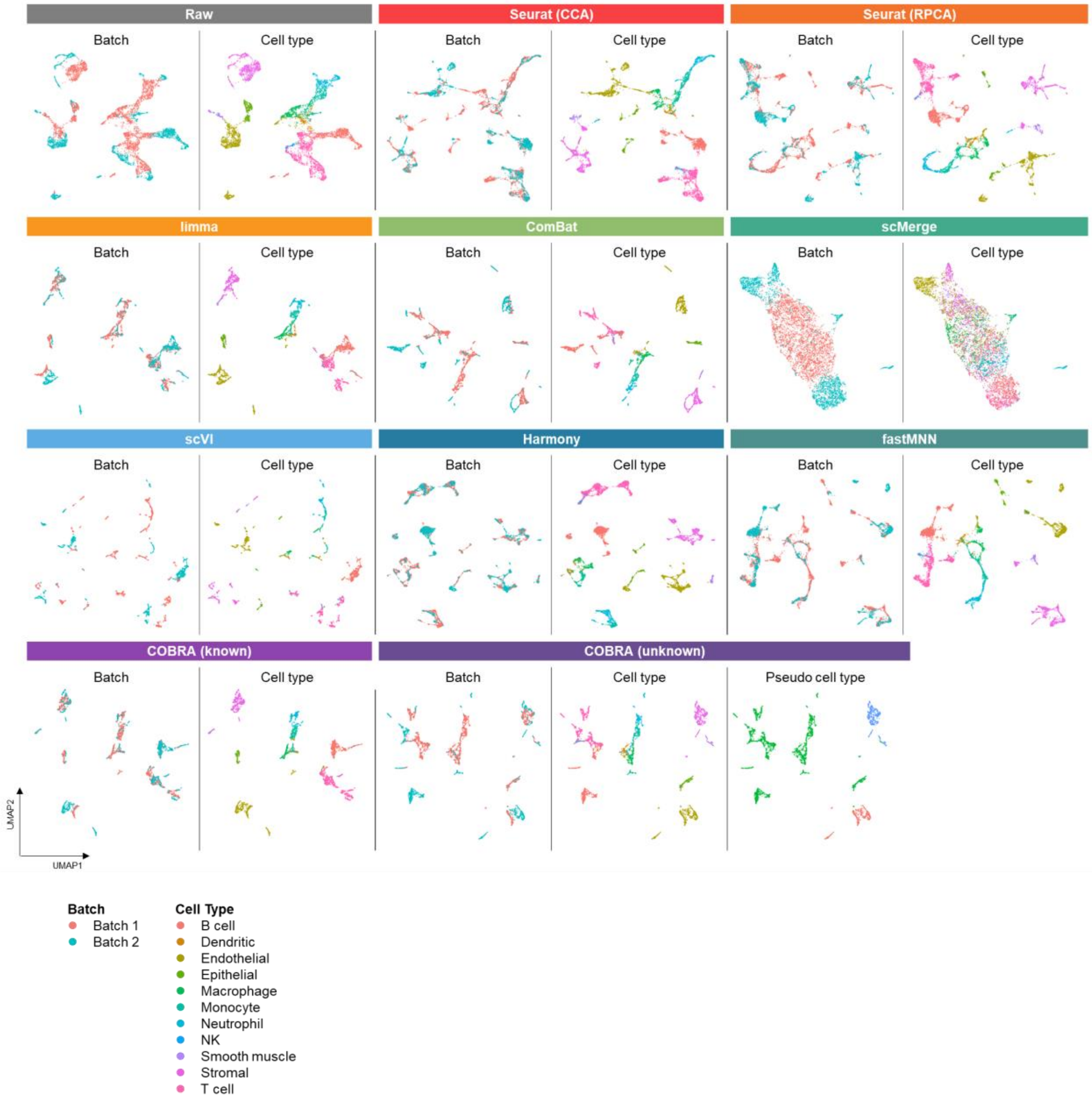

**Supplementary Figure S6. UMAP visualization for human pancreas dataset**

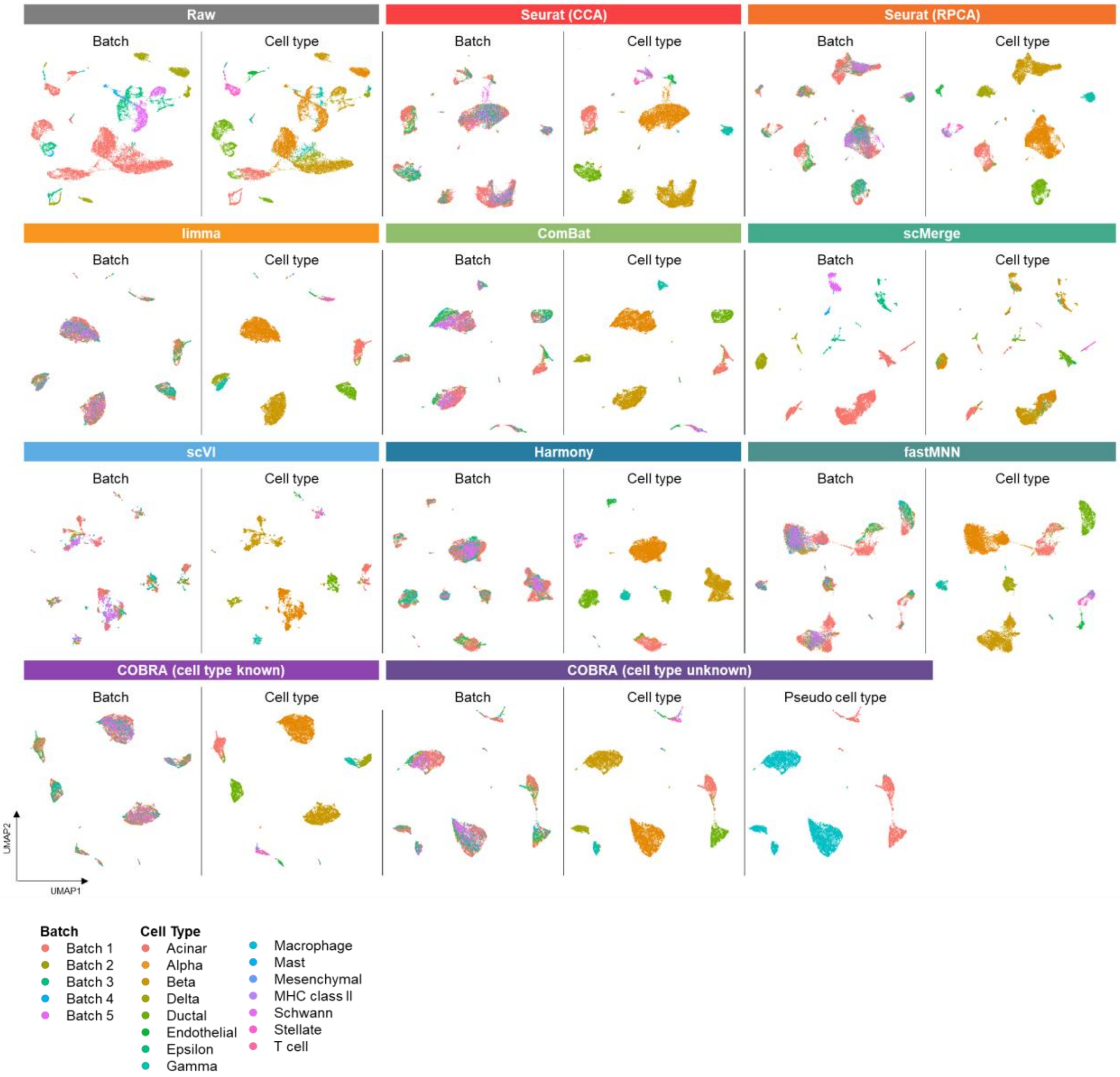

#### Supplementary Figure S7. UMAP visualization for human PBMC dataset

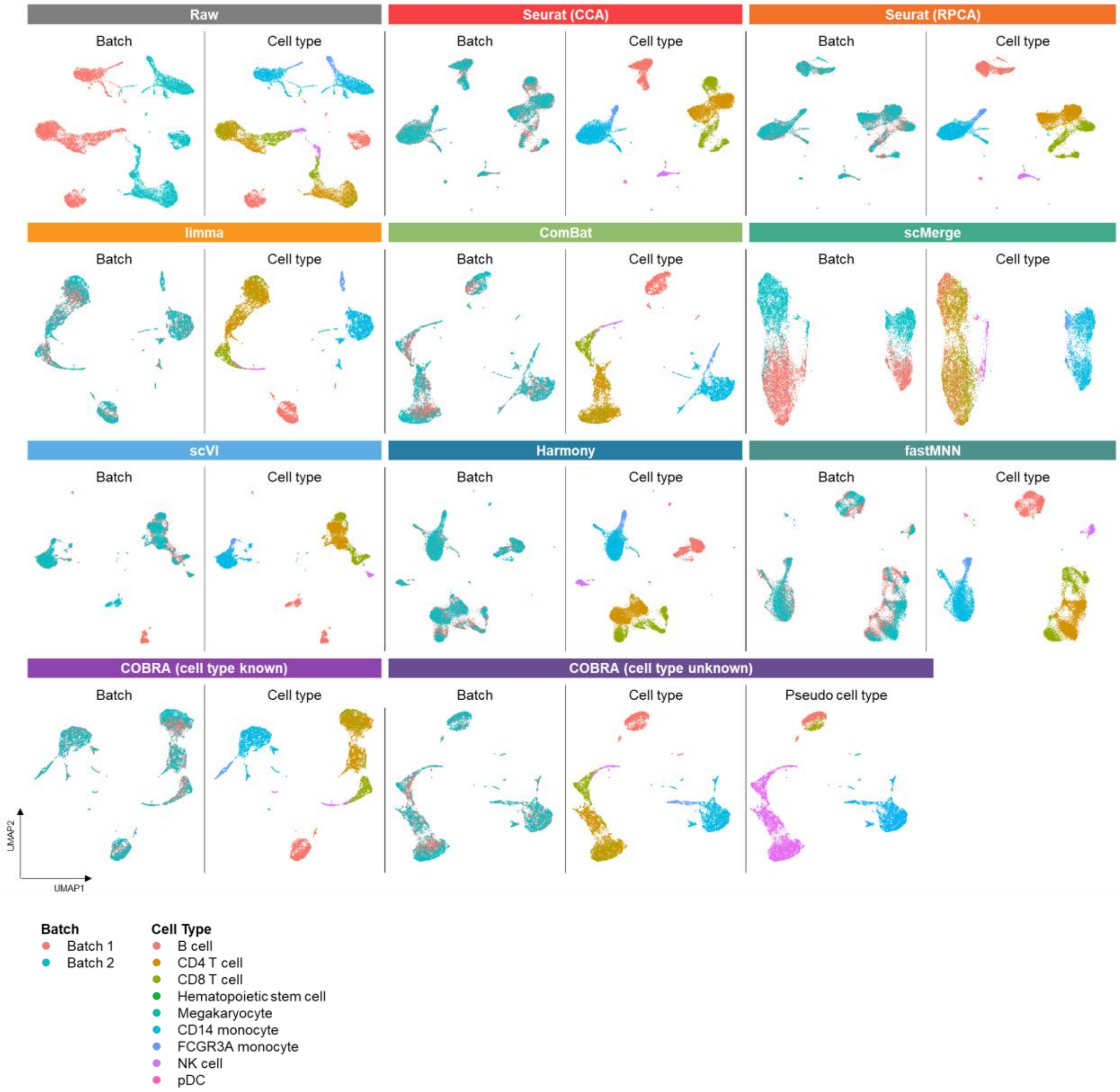

### Supplementary Figure S8. UMAP visualization for mouse retina

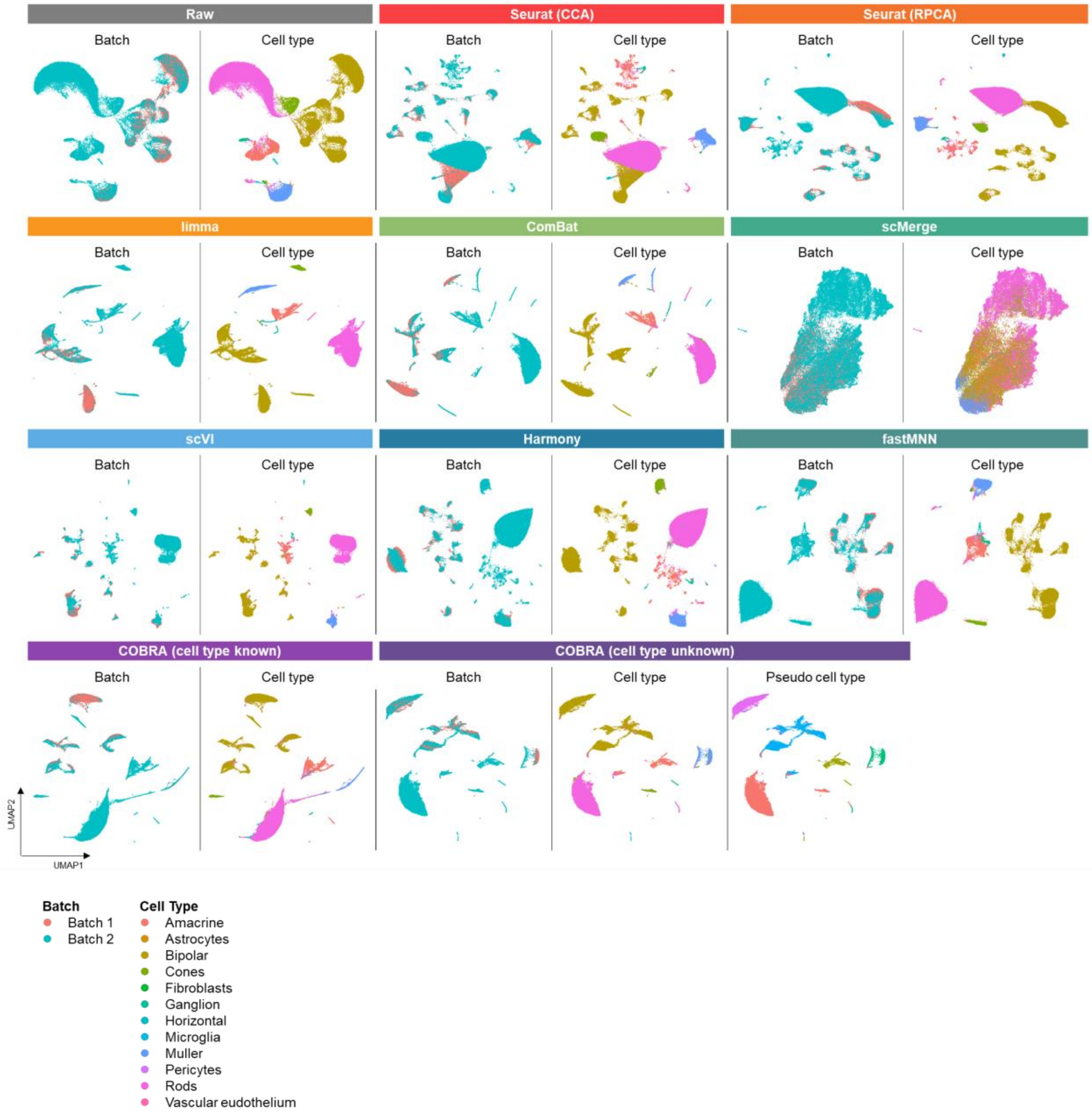

**Supplementary Figure S9. UMAP visualization for mouse hematopoietic stem and progenitor cells dataset**

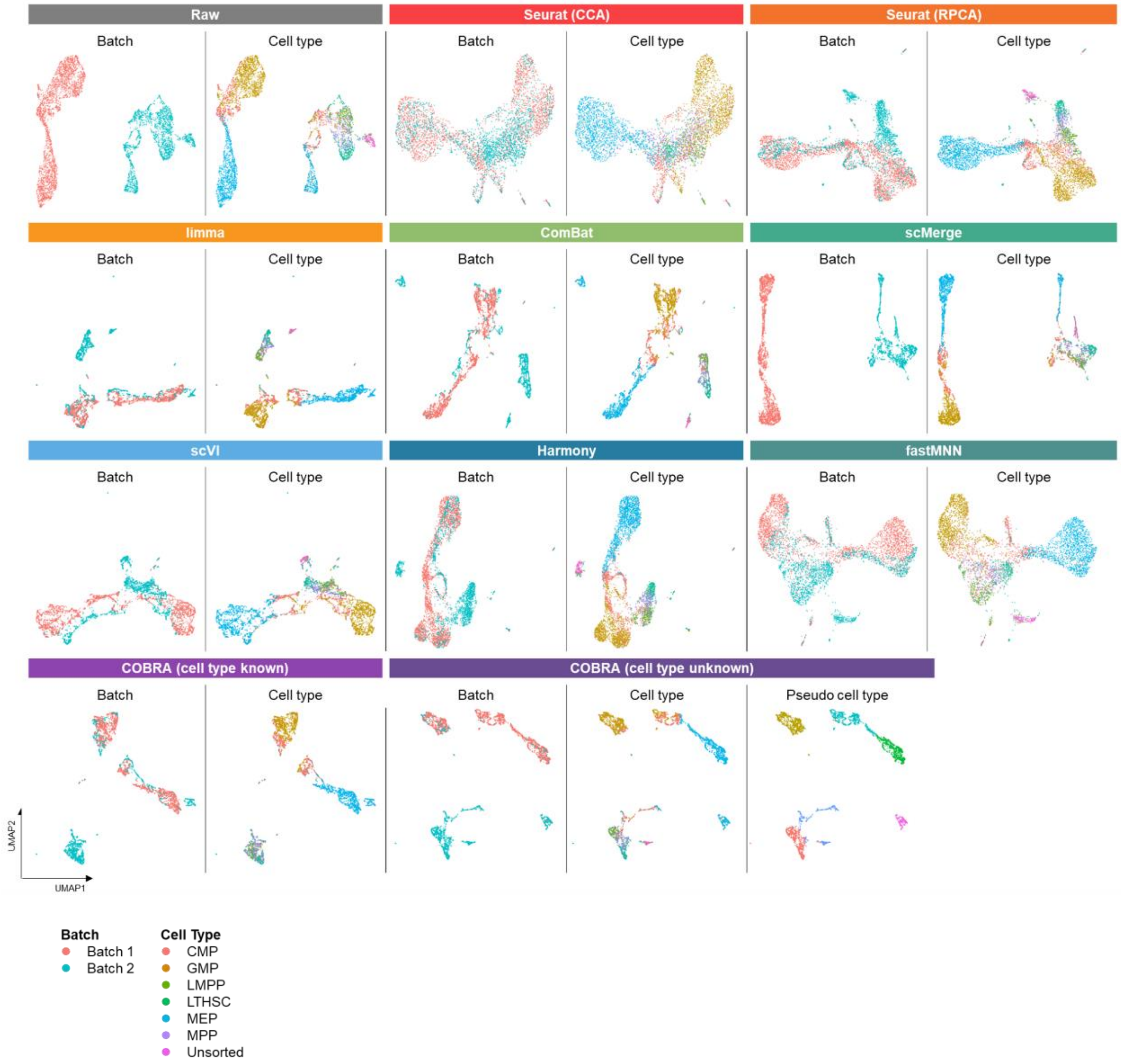

#### Supplementary Figure S10. UMAP visualization for mouse brain dataset

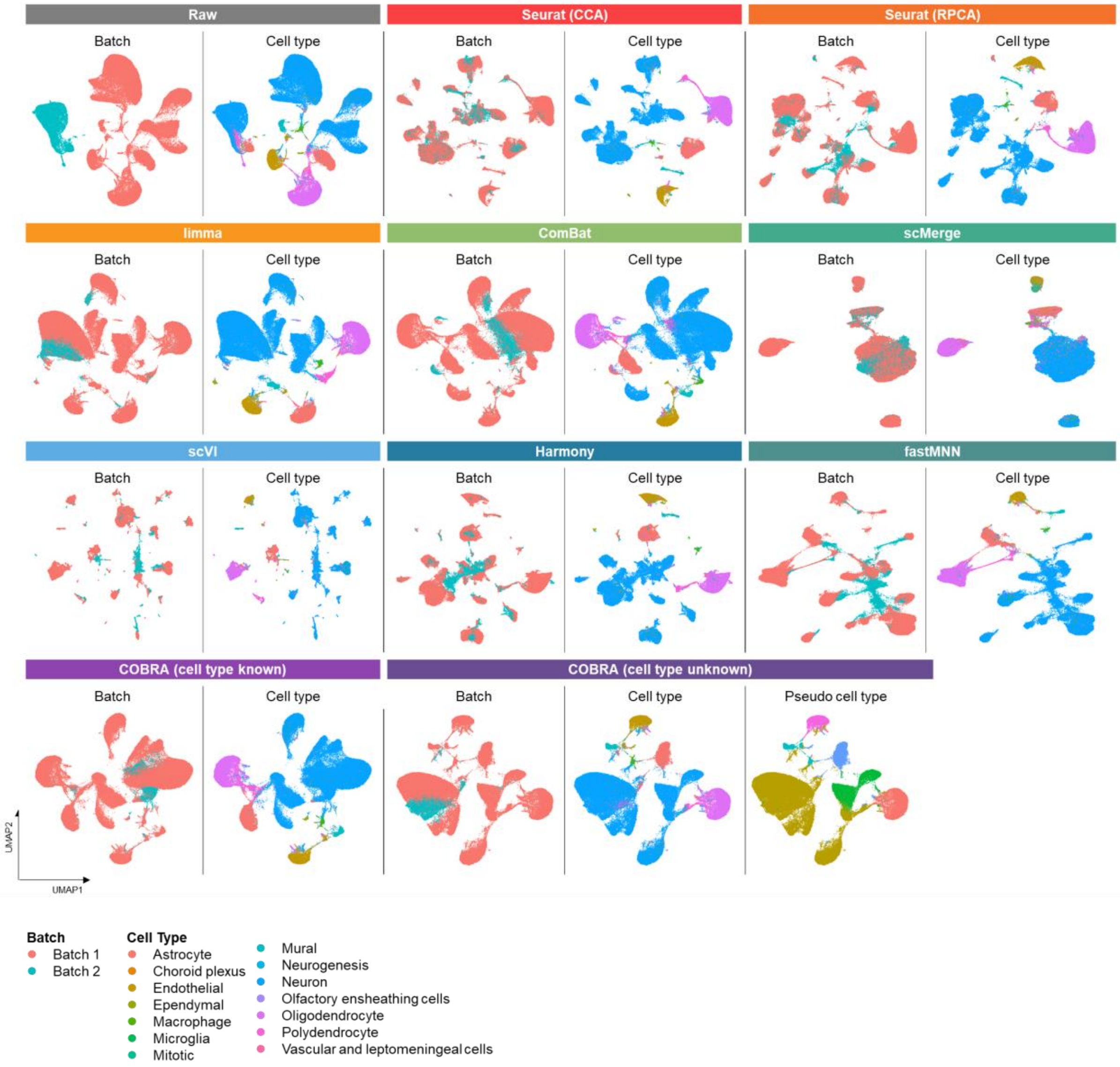
